## Supplementary material for "The Trouble Interpreting fMRI Studies in Populations with Cerebrovascular Risk: The Use of a Subject-Specific Hemodynamic Response Function in a Study of Age, Vascular Risk, and Memory": Matlab Code for Converting HRF

```
%   Create new GLM analysis using a provided subject-specific HRF
%   after an analysis has already been conducted using the canonical HRF
%  
%  Usage: cHRF_to_sHRF(hrf_csv, P, [outdir])
%
%  Where:
%
%	hrf_csv: The path to a csv file with the first column containing the folder name of
%	each subject (e.g., S_001) and the remaining columns representing the
%	custom HRF (e.g., taken from a Finite Impulse Response analysis from a
%	different task). The file should contain no headers.
%
%   P:   A text file of containing the path names to each GLM directory for 
%   each subject included in the analysis (e.g.,
%   /Path/Subject/GLM_cHRF_dir/)
%
%	outdir (optional):	Name of the new directory that will be created with 
%   the sHRF.  Default is 'GLM_sHRF'.

%
%  e.g.:
%     hrf_csv = '/Path/FIR.csv';
%     P = '/Path/Participant_List.txt';
%     cHRF_to_sHRF(hrf_csv,P);    % default outdir
%
%
%  - Ian McDonough
%
function cHRF_to_sHRF(varargin)
   
    if nargin < 1 || isempty(varargin{1})
        warning('Not enough inputs provided');
        return
    end
    
    if ~isempty(varargin{1})
        hrf_csv = varargin{1};
    else
        hrf_csv = spm_select(1,'file','Select hrf csv');
    end
    
    if ~isempty(varargin{2})
        P = varargin{2};
    end
    
    if ~isempty(varargin{3})
        outdir = varargin{3};
    end
    
    if nargin >= 1 && isempty(varargin{2})
        P = spm_select(inf,'dir','Select 1st level GLM directories');
        args = cellstr(P);
    end

    if nargin >= 1 && isempty(varargin{3})
        outdir = 'GLM_sHRF';
    end

    try 
        %Load subject-specific HRF
        hrf = readtable(deblank(hrf_csv));
    catch ME
        rethrow(ME)
    end
    
    if ~exist('args','var')
        try
            %Load Participant List
            fileID = fopen(deblank(P),'r');
            formatSpec = '%s%[^\n\r]';
            delimiter = '';
            dataArray = textscan(fileID, formatSpec, 'Delimiter', delimiter,  'ReturnOnError', false);
            fclose(fileID);
            raw = repmat({''},length(dataArray{1}),length(dataArray)-1);
            for col=1:length(dataArray)-1
                raw(1:length(dataArray{col}),col) = dataArray{col};
            end
            numericData = NaN(size(dataArray{1},1),size(dataArray,2));
            rawNumericColumns = {};
            rawCellColumns = raw(:, 1);
            args = raw;
            %% Clear temporary variables
            clearvars P delimiter formatSpec fileID dataArray ans raw col numericData rawNumericColumns rawCellColumns;
        catch ME
            rethrow(ME)
        end
    end

    for j = 1:size(args, 1) % Loop through each subject
        
        if ~exist(args{j,1},'dir')
            warning('%s does not exist',args{j,1});
            continue
        end
        
        cd(args{j,1})
        
        %If trying to re-run, first remove directory
        if exist(fullfile('../',outdir),'dir')
            rmdir(fullfile('../',outdir),'s');
%             rmdir('../*sHRF_s8*','s')
        end
        
        if exist(fullfile('../',outdir,'beta_0001.nii'),'file')
            warning('New HRF already calculated for %s',args{j,1})
            continue
        else
            mkdir(fullfile('../',outdir))
            try
                copyfile('SPM.mat',fullfile('../',outdir,'SPM.mat'));
            catch ME
                display(ME.message)
                continue
            end

            cd(fullfile('..',outdir))
            
            pathparts = explode(pwd,'/');
            subj = pathparts{1,end-1};
            
            try
                %find subject's HRF
                sHRF = table2array(hrf(find(strcmp(subj, hrf{:,1})),2:end));
            catch ME
                display(ME.message)
                continue
            end

            % load SPM.mat specification with canonical HRF
            load('SPM.mat')

            dt = SPM.xBF.dt;
            res   = SPM.xBF.T;
            
            try
                lenh = length(sHRF);
                sHRF = interp1((1:lenh)', sHRF, (1:(1/res):lenh), 'linear', 'extrap');
                sHRF = sHRF ./ sum(sHRF);
            catch ME
                display(ME.message)
                continue
            end
                

            SPM.xBF.bf = sHRF';
            if SPM.xBF.order > 1
                order_orig = SPM.xBF.order;
                SPM.xBF.order = 1;
            end
            SPM.xBF.name = 'hrf';
            SPM.swd = pwd;
            
            len_sess = length(SPM.Sess);
            Num_Con = [];
            Col_Remove = [];
            for k=1:len_sess
                tmp = 1;
                len_fc = length(SPM.Sess(k).Fc);
                for l=1:len_fc
                   SPM.Sess(k).Fc(1,l).i = tmp;
                   SPM.Sess(k).Fc(1,l).p = 1;
                   tmp=tmp+1;
                end
                if k == 1
                    col_length=(length(SPM.Sess(k).col)-(length(SPM.Sess(k).U)*(order_orig-1)));
                    SPM.Sess(k).col = [1:1:col_length];
                    tmp_end = col_length;
                else
                    col_length=(length(SPM.Sess(k).col)-(length(SPM.Sess(k).U)*(order_orig-1)));
                    SPM.Sess(k).col = [tmp_end+1:1:col_length+tmp_end];
                    tmp_end = length(SPM.Sess(k).col)-(length(SPM.Sess(k).U)*(order_orig-1));
                end
                Num_Con = [Num_Con; length(SPM.Sess(k).U)];
            end
            for k=1:len_sess
                con_tmp = 1;
                if order_orig == 2
                    if k == 1
                        for l = 1:length(SPM.Sess(k).U)
                            Col_Remove = [Col_Remove (con_tmp+1)];
                            con_tmp = Col_Remove(1,end)+1;
                        end
                    else
                        new_Col = Col_Remove +  Num_Con(k,1)*order_orig + size(SPM.Sess(k).C.C,2);
                        Col_Remove = [Col_Remove new_Col];
                    end
                elseif order_orig == 3
                    if k == 1
                        for l = 1:length(SPM.Sess(k).U)
                            Col_Remove = [Col_Remove (con_tmp+1) (con_tmp+2)];
                            con_tmp = Col_Remove(1,end)+1;
                        end
                    else
                        new_Col = Col_Remove +  Num_Con(k,1)*order_orig + size(SPM.Sess(k-1).C.C,2);
                        Col_Remove = [Col_Remove new_Col];
                    end
                end
            end
            
            % Remove Derivatives from Design Matrix
            SPM.xX.X(:,Col_Remove) = [];
            SPM.xX.name(:,Col_Remove) = [];
            SPM.xX.pKX(Col_Remove',:) = [];
            SPM.xX.Bcov(:,Col_Remove) = [];
            SPM.xX.Bcov(Col_Remove',:) = [];
            SPM.xX.nKX(:,Col_Remove) = [];
            SPM.xX.xKXs.X(:,Col_Remove) = [];
            SPM.xX.xKXs.ds(Col_Remove,:) = [];
            SPM.xX.xKXs.u(:,Col_Remove) = [];
            SPM.xX.xKXs.v(:,Col_Remove) = [];
            SPM.xX.xKXs.v(Col_Remove,:) = [];
            SPM.xX.xKXs.rk = length(SPM.xX.xKXs.v);
            SPM.xX.iC = 1:1:(size(SPM.xX.X,2)-2);
            SPM.xX.iB = [(size(SPM.xX.X,2)-1) (size(SPM.xX.X,2))];
            SPM.xX.trRV = SPM.xX.trRV + sum(Num_Con)*(order_orig-1);
            SPM.xX.trRVRV = SPM.xX.trRVRV + sum(Num_Con)*(order_orig-1);
            SPM.xX.erdf = SPM.xX.erdf + sum(Num_Con)*(order_orig-1);
            SPM.xsDes.Basis_functions = 'sHRF';
          
            
            SPM.Vbeta(:,Col_Remove) = [];
            for k=1:length(SPM.Vbeta)
                SPM.Vbeta(1,k).fname = sprintf('beta_%05d.nii',k);
            end
            
            SPM = rmfield(SPM, 'xVol');
            SPM = rmfield(SPM, 'xCon');
         
            
            save('SPM.mat','SPM')
            % Estimate new SPM.mat
            spm_spm(SPM);
          
            % Create new contrasts
            matlabbatch = [];
            matlabbatch{1}.spm.stats.con.spmmat = cellstr(strcat(pwd,'/SPM.mat'));
            matlabbatch{1}.spm.stats.con.consess{1}.tcon.name = 'Correct > Baseline';
            matlabbatch{1}.spm.stats.con.consess{1}.tcon.convec = [1 0];
            matlabbatch{1}.spm.stats.con.consess{1}.tcon.sessrep = 'replsc';
            matlabbatch{1}.spm.stats.con.consess{2}.tcon.name = 'Incorrect > Baseline';
            matlabbatch{1}.spm.stats.con.consess{2}.tcon.convec = [0 1];
            matlabbatch{1}.spm.stats.con.consess{2}.tcon.sessrep = 'replsc';
            matlabbatch{1}.spm.stats.con.consess{3}.tcon.name = 'Correct > SM_Incorrect';
            matlabbatch{1}.spm.stats.con.consess{3}.tcon.convec = [1 -1];
            matlabbatch{1}.spm.stats.con.consess{3}.tcon.sessrep = 'replsc';
            matlabbatch{1}.spm.stats.con.consess{4}.tcon.name = 'Incorrect > SM_Correct';
            matlabbatch{1}.spm.stats.con.consess{4}.tcon.convec = [-1 1];
            matlabbatch{1}.spm.stats.con.consess{4}.tcon.sessrep = 'replsc';
            matlabbatch{1}.spm.stats.con.delete = 1;
            
            try
                spm_jobman('run',matlabbatch);
            catch ME
                display(ME.message)
                continue
            end
        end
        display(sprintf('Done with %s', subj));
        clearvars -except P args j outdir hrf
    end
    
return;
```
